# Atypical somatosensory adaptation in adults on the autism spectrum: a high-density electrophysiological (EEG) mapping study

**DOI:** 10.64898/2026.01.05.697771

**Authors:** Emily L. Isenstein, Eve R. Lang, Edward G. Freedman, John J. Foxe

**Author notes:** Co-Senior Authorship (Edward G. Freedman, John J. Foxe).

## Abstract

Adaptation to repetitive sensory inputs promotes efficient neural processing by attenuating responses to redundant information and reallocating resources to novel stimuli. Reduced adaptation has been proposed to contribute to atypical sensory reactivity in autism, but the physiological mechanisms underlying tactile adaptation remain poorly understood. Here, we examined short-term adaptation to repetitive vibrotactile stimulation in autistic and neurotypical adults using high-density electrophysiological recordings. Fifty participants (18-44 years; 25 autistic; 25 neurotypical), received sequences of four brief vibrations to the index fingertip while viewing silent videos. Neural responses were analyzed for an early negative deflection (N1, ∼100 milliseconds) indexing basic stimulus recognition, and a later positive deflection (P2, ∼200-300 milliseconds) indexing higher-order contextual and attentional processing. Adaptation was quantified as changes in response magnitude across the four vibrations. The N1 did not differ between groups, showing minimal change across repetitions, indicating comparable processing of basic tactile features. In contrast, the P2 was significantly larger overall in the autistic group. Across both groups, responses to the first vibration in each sequence were greater than responses to subsequent vibrations, reflecting re-sensitization following the inter-sequence interval. Autistic participants exhibited consistently amplified P2 responses to initial vibrations, suggesting heightened re-sensitization rather than impaired within-sequence adaptation. Associations between neural responses and clinical measures of autistic traits and tactile sensitivity were modest. These findings indicate that autistic adults show amplified higher-order neural responses to tactile input alongside preserved short-term adaptation. Heightened re-sensitization to repeated touch may reflect shortened refractory periods, contributing to sensory hyper-reactivity and increased perceptual load.

## Introduction

Adaptation to repetitive sensory stimuli allows for progressive reduction of the magnitude of neural responses, and is understood as a mechanism to divert cortical and attentional resources away from the processing of redundant sensory information (Andrade et al., 2015; Groves & Thompson, 1970; Harris, 1943; Megela & Teyler, 1979; Müller et al., 1999; Thompson & Spencer, 1966), potentially freeing up valuable perceptual and cognitive resources to be allocated to novel, more salient information (Eisenstein et al., 2001; Rankin et al., 2009). Lack of adaptation could potentially lead to dysregulation of sensory-perceptual processing, creating an imbalance in the relative salience of novel and repeated sensory information (Andrade et al., 2016; Francisco et al., 2022; Podoly & Ben-Sasson, 2020; Schwartz et al., 2004).

Atypical adaptation has been reported in autism spectrum disorder (ASD) across the sensory systems. For example, anomalous adaptation to tactile inputs has been reported behaviorally (Puts et al., 2014; Sapey-Triomphe et al., 2023) and in neuroimaging work (Green et al., 2015; Puts et al., 2017), consistent with findings in the auditory and visual domains (Barry & James, 1988; Dwyer et al., 2023; Gandhi et al., 2021; Jamal et al., 2021; Lawson et al., 2015; Pellicano et al., 2007). These observations are part of a wider pattern of atypical responses to tactile information found across the autism spectrum (American Psychiatric Association, 2013; Baranek et al., 1997; Hashimoto et al., 1986; Rogers et al., 2003). For example, while threshold detection of light touch is largely equivalent between neurotypical and autistic individuals (Cascio et al., 2008; Güçlü et al., 2007; O’Riordan & Passetti, 2006), responses to vibrotactile stimuli have been shown to be dysregulated in autism (Blakemore et al., 2006; Mikkelsen et al., 2018; Puts et al., 2014; Russo et al., 2010; Tavassoli et al., 2016). However, the underlying physiological properties that drive these differences remain poorly characterized. A proposed mechanism of reduced sensory adaptation in autism may underlie this broadly dysregulated sensory profile (Guiraud et al., 2011; Sinha et al., 2014; Webb et al., 2010). Under this theory, decreased suppression of redundant sensory input could translate to heightened responses to incoming stimuli that can burden, or even overwhelm, the sensory-processing system. An inability to cope with this extra environmental information may result in hyper-reactivity to sensations, whereas exhaustion of the system may result in hypo-reactivity to inputs or a “shutdown”. Indeed, sensitivity to sensory inputs has been shown to predict discrimination of tactile inputs presented after an adapting stimulus (McKernan et al., 2020).

Here, we focus specifically on tactile adaptation in ASD as atypical behavioral reactivity to somatosensory inputs is very commonly reported in this population and has significant implications for quality of life (Ferrer Knight & Birtles, 2025; Foss-Feig et al., 2012; Isenstein et al., 2024; Jamioł-Milc et al., 2021; Mammen et al., 2015). Tactile inputs that offer consistent sensory feedback - such as wearing a shirt, sitting on a chair, or holding someone’s hand - typically follow a standard course of adaptation that has been demonstrated behaviorally and electrophysiologically using somatosensory evoked potentials (SEPs) (Angel et al., 1985; Nakajima & Imamura, 2000; van der Miesen et al., 2024; Yamaguchi & Knight, 1991). As sensory adaptation mechanisms evolve over the course of physiologic maturation (Freedman et al., 1987; Muenssinger et al., 2013; Zanini et al., 2016), atypical patterns of development may influence the trajectory of tactile adaptation capacities. For example, individuals with Tourette syndrome (Puts et al., 2015), fibromyalgia (Montoya et al., 2006), cerebral palsy (Kurz et al., 2017), and schizophrenia (Andrade et al., 2016) have been shown to have dysregulated adaptation to tactile stimuli.

A deeper understanding of adaptation mechanisms in autism is required to further assess how these dysregulated processes may impact higher-order sensory-perceptual processes. Yet to our knowledge, adaptation to tactile inputs has not yet been tested electrophysiologically in idiopathic autism. High-density electrophysiology (EEG) offers the benefit of a rapid, non-invasive, and objective measure of neurological processing that is accessible to individuals of all ages and cognitive abilities, allowing for the extension of adaptation findings already conducted in behavioral and neuroimaging work. Leveraging the high temporal resolution of the EEG technique, we assessed the time-course and extent of sensory habituation in the tactile domain, focusing on the well-characterized N1 and the P220 electrophysiological components. The N1 component is thought to reflect basic recognition of stimulus properties (Cauller, 1995; Desmedt et al., 1976; Wang et al., 2008), whereas a positive peak 200-300 ms post-stimulus is thought to index higher-order processing responsible for attributing context, relevance, and attention to the input (Liu et al., 2021; Schröder et al., 2021; Yamaguchi & Knight, 1991).

Here, we examined the short-term adaptation of vibrotactile stimuli in groups of autistic and neurotypical adults. By repeatedly presenting sequences of four vibrations to the index fingertip, we captured adaptation of neurophysiological responses to somatosensory stimuli on the order of milliseconds. Based on our prior work (Isenstein et al., 2024; Isenstein et al., 2025), we predicted that early stimulus recognition processes reflected by the N1 would be somewhat reduced in the autism group but with overall comparable adaptation when compared to neurotypical peers. We further hypothesized that later higher-order attentional salience processes reflected by the P220 would show increased magnitude activity and decreased adaptation over time in Autism, reflecting a reduced ability to divert attentional resources away from redundant tactile information.

## Methods

### Participants

Fifty adults between the ages of 18 and 45 were recruited, 25 had a diagnosis of autism and 25 were neurotypical controls. All reported normal hearing and normal or corrected-to-normal vision. None reported a history of traumatic brain injury or psychosis-related disorder. Both groups were recruited from the local Rochester New York area. Members of the control group (mean age: 24.77, StdErr: 1.32, 11M/14F) did not have first-degree biological relatives with an autism diagnosis. The autism group (mean age: 24.70, StdErr: 1.44, 13M/12F) were all previously diagnosed with autism by a clinician and met criteria on the Autism Diagnostic Observation Schedule 2 (ADOS-2) (Lord et al., 2012) at the time of study participation. The Wechsler Abbreviated Scale of Intelligence (Wechsler, 2011) was used to determine the IQs of all participants, but there was no IQ cutoff used as exclusion criterion for participation. The average IQ of the control group was 110.17 (StdErr: 2.74, one participant’s data were unavailable due to administration error) and the average IQ of the autism group was 110.72 (StdErr: 3.10). No participants took medication for a psychosis-related condition. Participants in the autism group who took stimulant medication (n = 7) withheld these medications on the day of the EEG; no participants in the neurotypical group took stimulant medication. Study participants or guardians, as appropriate, completed the Social Responsiveness Scale 2 (Adult or Adult Self-Report; SRS-2) (Constantino & Gruber, 2012), the Adolescent/Adult Sensory Profile (SP) (Brown & Dunn, 2002) and the Adult ADHD Self-Report Screening Scale for DSM-5 (ASRS-5) (Ustun et al., 2017). See Table 1 for full demographic information.

**Table 1.** Demographic data.

|  | NT | ASD | *p* - value |
| --- | --- | --- | --- |
| Age(SE) | 24.77(1.32) | 24.70(1.44) | 0.85 |
| Sex | 11M/14F | 13M/12F | 0.58 |
| Gender |  |  |  |
| Man | 11 | 13 |  |
| Woman | 13 | 8 |  |
| Nonbinary | 1 | 4 |  |
| Handedness |  |  |  |
| *Right* | 23 | 23 |  |
| *Left* | 1 | 0 |  |
| *Ambidextrous* | 0 | 1 |  |
| *Not Reported* | 2 | 2 |  |
| *Race* |  |  |  |
| *White* | 12 | 18 |  |
| *Black or African American* | 3 | 1 |  |
| *Asian* | 5 | 3 |  |
| *Hawaiian or Pacific Islander* | 0 | 0 |  |
| *American Indian* | 0 | 0 |  |
| *Multiple Races\** | 3 | 1 |  |
| *Unknown/did not report* | 2 | 1 |  |
| *Ethnicity* |  |  |  |
| *Hispanic/Latino* | 5 | 1 |  |
| *Not Hispanic/Latino* | 18 | 22 |  |
| *Unknown/refused to answer* | 2 | 2 |  |
| *WASI Full Scale* | 109.82 (SE = 2.89) | 111.54 (3.28) | 0.70 |
| *ASRS-5* | 7.75 (0.72) | 12.88 (0.87) | $4.20 \times 10^{-5}$ |
| *SRS-2 Total Score* | 35.52 (3.46) | 95.28 (5.93) | $1.96 \times 10^{-11}$ |
| *SP Touch Subscale* | 29.04 (1.13) | 33.72 (1.62) | 0.02 |

\*Mixed race participants reported being: Black & White, Mestizo, American Indian & White, or decline to provide further detail.

Participants in the autism group were informally asked about their preference regarding person first (person with autism) or identity first (autistic person) language. There was no clear majority regarding preference; accordingly, person and identity first language are both used throughout this paper.

All participants or their legal guardians gave written informed consent, and additional verbal and written assent was obtained as appropriate. All procedures were reviewed and approved by the Research Subjects Review Board (RSRB) at the University of Rochester (STUDY00002036) and conformed with the tenets for ethical conduct of human subjects’ research laid out in the Helsinki declaration (“World Medical Association Declaration of Helsinki: ethical principles for medical research involving human subjects,” 2013). Participants received an hourly rate for their participation.

### EEG Stimuli and Task

An Adafruit 1201 vibrating mini-motor disc (Arduino, Turin, Italy) snuggly fastened around the right index fingertip was used to deliver vibrations. The mini-motor disc was controlled by an Arduino Uno microcontroller operated through Presentation® software (Version 18.0, Neurobehavioral Systems, Inc., Berkeley, CA). All vibrations were 5 V, 183.33 Hz, and 100 ms. Participants completed the experiment in an electrically-shielded and sound-attenuating EEG booth (IAC Acoustics, North Aurora, IL, USA) with earplugs (MPOW HM035A; Longgang, Guangdong, China) to reduce air-conducted noise. Participants were told to ignore the vibrations and watched a silent video of their choosing as a distraction. They were instructed to position their hand with palm facing upward with the mini-motor disc off the table so the vibration would not be affected. Vibrations were presented in sequences of four, with a stimulus onset asynchrony (SOA) of 500 ms; each sequence was separated by an inter-stimulus interval (ISI) of 4000 ms. A total of 200 sequences of four were presented over approximately 20 minutes. All participants completed this experiment following a short break after completing a separate EEG experiment lasting 1-1.5 hours, and electrode impedance was checked and corrected as needed prior to beginning the present experiment.

A 128-channel BioSemi EEG system (Biosemi ActiveTwo, Amsterdam, The Netherlands) was used to record continuous high-density EEG at 512 Hz with the default Biosemi decimation filter of 1/5 of this rate. The EEGLAB toolbox (version 14.1.2; Delorme & Makeig, 2004) with MATLAB 2021a software (Mathworks, Natick, MA, USA) was used for preprocessing. Data were filtered from 0.01 to 40 Hz and outlier channel with amplitudes ± 3 standard deviations from the mean were interpolated. The autism group had on average 14.48 channels removed, and the neurotypical group had on average 15.44 channels removed. Data were re-referenced to the average reference and automatic artifact rejection was used. Data were epoched from -500 to 2000 ms around the onset of the first vibration of the sequence, and baseline corrected to the 100 ms prior to the sequence. Trials containing amplitudes +/- 200 µV were rejected. The autism group had an average of 25.28 trials removed (SD: 24.34), and the neurotypical group had an average of 18.40 trials removed (SD: 15.81)

### A Priori Analyses

For N1 amplitude, the average of a cluster of 5 electrodes centered in the negative polarity region of the associated topography plot was selected to maximize the N1 signal (Figure 2C). For P220 amplitude, the average of a cluster of 5 electrodes centered on the Cz vertex (Figure 2C) were defined as the region of interest based on prior adaptation studies (Isenstein et al., 2022; Mancini et al., 2018; Rosburg et al., 2010).

**Figure 1.**
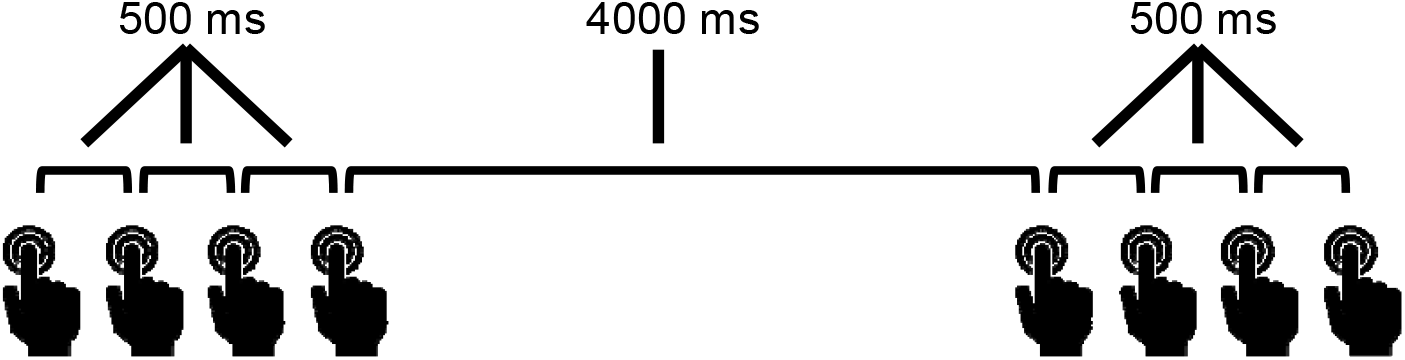
A visual representation of the somatosensory paradigm.

**Figure 2.**
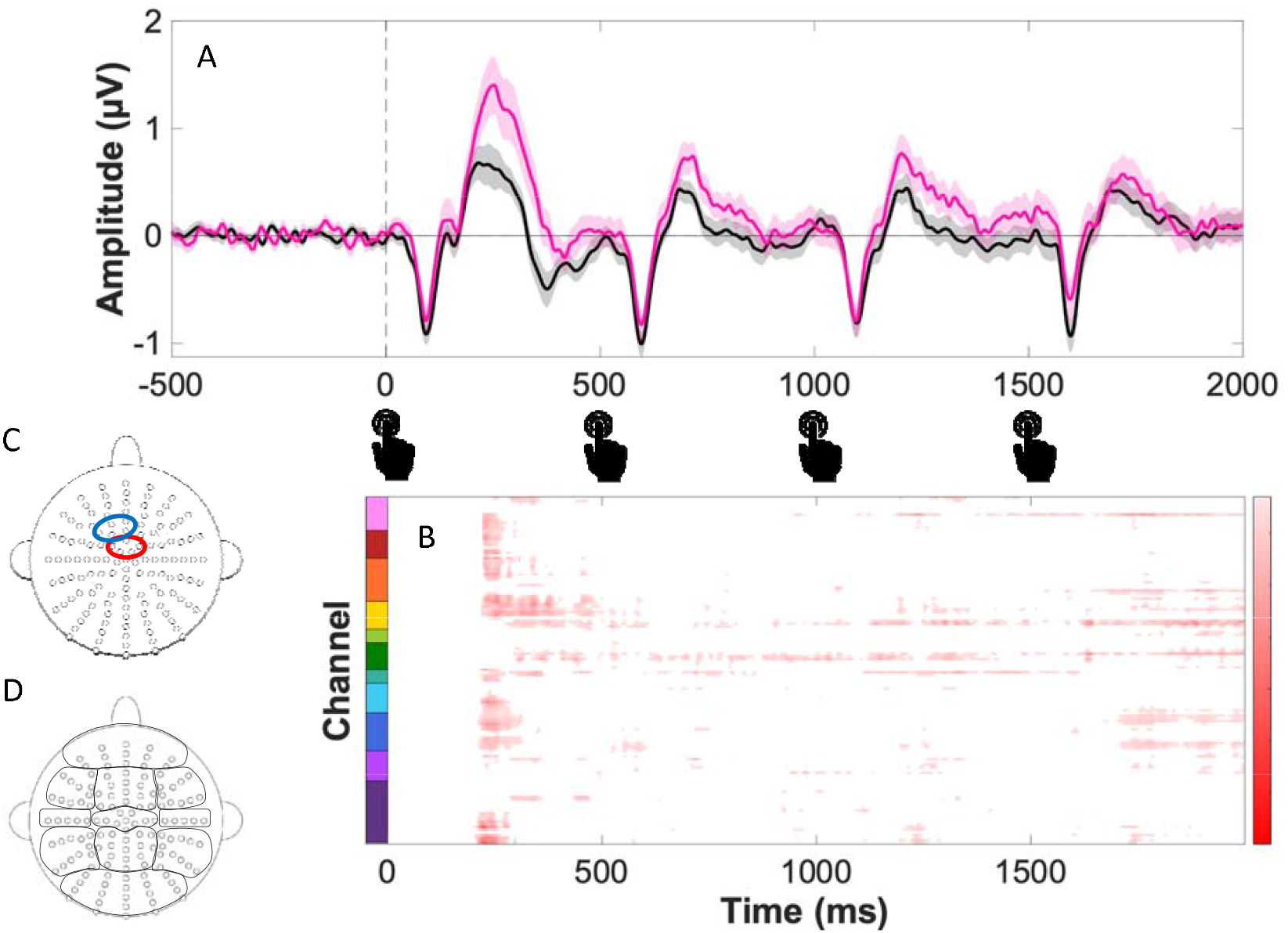
(A) Grand average ERPs for the ASD (pink) and NT (black) for the entire trial duration. (B) Statistical cluster plot comparing difference between ASD and NT ERPs represented using p-values (p <0.05 in red, p > 0.05 in white), red shading indicates statistically significant difference in μV. (C) Scalp plot of the 128-electrode array. The N1 electrode cluster is circled in blue and the P220 electrode cluster is circled in red. (D) The y-axis of Figure 2B is plotted on the 128-electrode array to demonstrate the physical location of each comparison.

**Figure 3.**
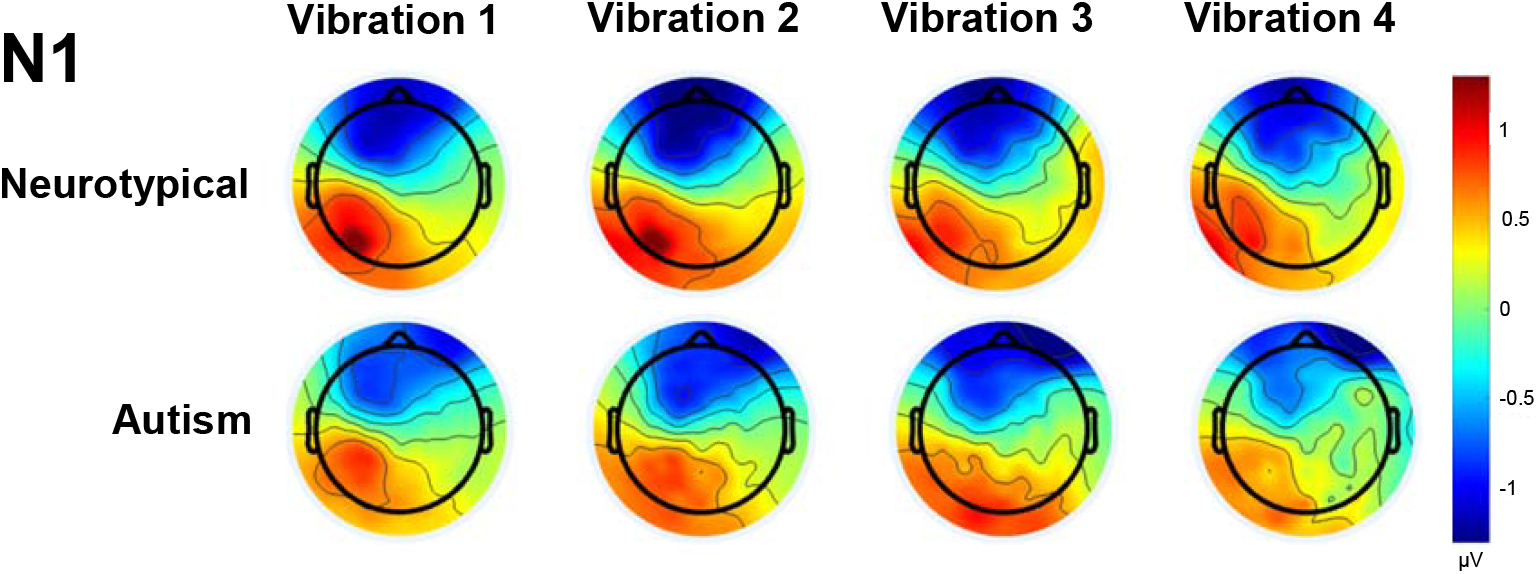
Scalp topographic maps of the N1 are shown for each group, representing the period from 95 - 105 ms post vibration.

**Figure 4.**
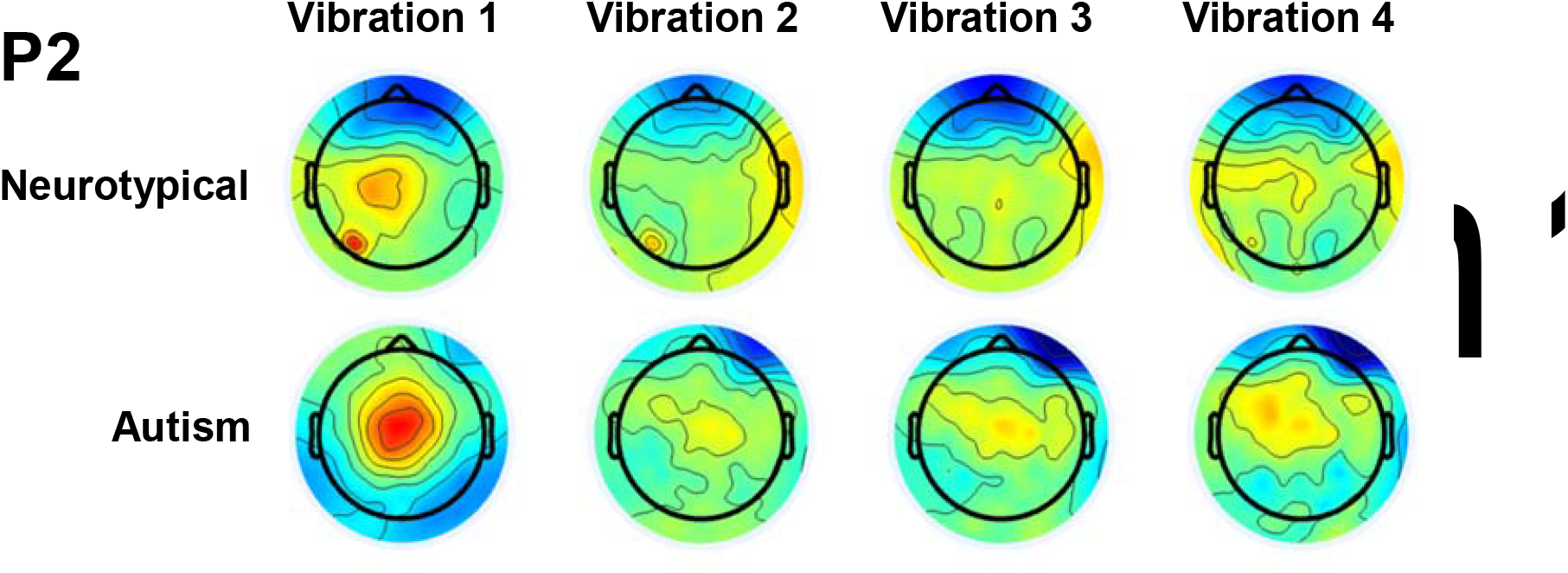
Scalp topographic maps of the P2 are shown for each group, representing the period from 190 - 240 ms post vibration.

Latency for the maximal negative peak for the N1 component was calculated as the maximum negative-going amplitude between 95-105 ms after each vibration across both groups. The same process was repeated for the P220 component, with maximum positive-going amplitudes calculated between 190-240 ms after each of the four vibrations.

Mixed-design analysis of variance (ANOVA) was used to compare amplitudes of the N1 and P220 components using a within-subjects factor of VIBRATION NUMBER (One, Two, Three, Four) and a between-subjects factor of GROUP (Neurotypical (NT), Autism (ASD)). Protected pairwise t-tests were conducted on the individual variables contributing to significant ANOVA findings.

Correlations were conducted between Vibration One N1 and P220 amplitude and select clinical measures including the SRS-2 total score and SP Touch subscale. P-values were corrected for multiple comparisons using a Benjamini-Hochberg <u>False Discovery Rate</u> of 0.05 (Benjamini & Hochberg, 1995).

## Results

Basic demographic information was compared between groups to identify any potential underlying differences between the neurotypical and autism cohorts. Race and ethnicity data are also included. There were no group differences in Sex, Age, or Full-Scale IQ. Additional comparisons of clinical characterization data are presented in Table 1, including the SRS-2, SP, and ASRS-5. The autism group had significantly higher ASRS-5, SRS-2 Total Score, and SP Touch Subscale scores (p < 0.05).

### A Priori Results

#### N1 Amplitude

There was no main effect of VIBRATION NUMBER (F(3,144) = 2.20, *p* = 0.11, η^2^ = 0.04) or GROUP (F(1,48) = 0.07, *p* = 0.79, η^2^ = 0.01). There was no interaction between VIBRATION NUMBER and GROUP (F(3,144) = 0.62, *p* = 0.57, η^2^ = 0.01). Sphericity was violated (*p* = 2.34*10^-3^).

#### P220 Amplitude

There was a main effect of VIBRATION NUMBER (F(3,144) = 11.73, *p* = 3.78*10^-5^, η^2^ = 0.20) and GROUP (F(1,48) = 6.29, *p* = 0.02, η^2^ = 0.12). There was no interaction between VIBRATION NUMBER and GROUP (F(3,144) = 0.86, *p* = 0.42, η^2^ = 0.02). Sphericity was violated. Pairwise comparison also showed that the P2 amplitude of Vibration One (Mean: 1.29, Std Err: 0.16) was significantly larger than Vibration Two (Mean: 0.81, Std Err: 0.08; *p* = 2.28*10^-4^, C.I. = 0.24, 0.73), Vibration Three (Mean: 0.83, Std Err: 0.09; *p* = 5.61*10^-4^, C.I. = 0.21, 0.71), and Vibration Four (Mean: 0.73, Std Err: 0.09; *p* = 2.01*10^-4^, C.I. = 0.28, 0.84). All other p’s > 0.17.

Post-hoc pairwise comparison between each of the four Vibrations between groups yielded a significantly larger P2 amplitude in the Autism group than the Neurotypical group (Mean: 0.63, Std Err: 0.20) in response to Vibration One (*p* = 0.05, C.I. = -1.25, -0.01), Vibration Two (*p* = 0.01, C.I. = -0.74, -0.10), and Vibration Three (*p* = 0.02, C.I. = -0.78, -0.06). All other *p*’s > 0.12. See Table 2 for mean and standard error amplitude values.

**Table 2.**

| | Vibration One<br>$\mu$ (SE) | Vibration Two<br>$\mu$ (SE) | Vibration Three<br>$\mu$ (SE) | Vibration Four<br>$\mu$ (SE) |
| --- | --- | --- | --- | --- |
| Neurotypical | 0.98 (0.22) | 0.60 (0.11) | 0.62 (0.13) | 0.59 (0.13) |
| Autism | 1.61 (0.22) | 1.02 (0.11) | 1.04 (0.13) | 0.88 (0.13) |

### Correlations

There was a significant correlation between P220 amplitude and SRS-2 total score (*r* = 0.33, *p* = 0.02), but this effect did not survive correction for multiple comparisons. All other correlations were non-significant (*p*’s > 0.26).

## Discussion

Understanding the underlying mechanisms behind tactile processing is critical for characterizing the biological processes that determine sensory reactivity and overload in autism. Here, we examined responsiveness and adaptation to repetitive tactile information in adults on the autism spectrum and neurotypical peers. To our knowledge, this experiment is the first to explore short-term adaptation to somatosensory stimulation in autistic adults in this manner.

Our primary analysis revealed that the autism group had overall larger amplitude P220 complexes in response to early vibrotactile inputs. This finding is consistent with our previous work that showed greater amplitudes of positive components in the 200-300 ms range in response to somatosensory stimuli in adults with autism as compared to neurotypical peers (Isenstein et al., 2024; Isenstein et al., 2025). In addition, the sequence of the vibrations had a strong effect, with a consistently elevated response to the first vibration compared to the subsequent vibrations, regardless of group. This effect demonstrates consistent re-sensitization to the tactile inputs following the gap of time between sequences. However, in conjunction with the general group differences, there is a pattern of heightened re-sensitization in the autism group as compared to the neurotypical group, indicating a possible shortened refractory period. This effect likely parallels the characteristic sensory reactivity that is a core diagnostic criterion for autism diagnosis (American Psychiatric Association, 2013). The lack of interaction between group and vibration number suggests comparable adaptation over the course of the vibration sequence between groups, highlighting re-sensitization as the driving factor that distinguishes the groups on this shorter time scale.

There were no differences between groups or across the sequence of vibrations with respect to the amplitude of the N1, suggesting equivalent sensory-perceptual processing of basic stimulus properties. This result is consistent with previous findings that did not find early negative adaptation within the ERPs in children presented with repetitive vibrotactile stimuli, although in this case the central scalp region was not included and adaptation was not calculated for a mid-latency positive component (Espenhahn et al., 2021). Although one would typically expect some degree of adaptation of the N1 of any sensory modality in the refractory period following repetitive identical stimuli (Angel et al., 1985; Bruin et al., 2000; Budd et al., 1998), the relatively long stimulus onset asynchrony of 500 ms may have contributed to a lack of significant adaptation over the sequence (Uppal et al., 2016; Wang et al., 2008). The extended 4000 millisecond gaps between trial sequences may have also affected sensitization and adaptation of the N1. Moreso, while auditory and visual adaptation have been well-mapped electrophysiologically in humans, there is limited literature recording how somatosensory inputs are handled. For example, somatosensory processing is generally slower and less precise than auditory processing (Butler et al., 2012; Tarkka et al., 1992; Villalonga et al., 2021), leading to potential differences in how various sensory processes are handled. Additionally, predictable timing of tactile inputs can cause suppression of the N1 (Ren et al., 2025), which is relevant for our particular task.

These analyses build upon the theory that autism is a disorder of prediction. If people with autism are unable to appropriately predict and modulate brain responses to repeated stimuli, it may lead to excessive cortical processing being allocated to the processing of repeated stimuli. Consequently, there may be an alteration in how novel stimuli are handled, affecting how individuals interact with the world around them. In the context of our collective findings, autism may present with a shortened refractory period to tactile inputs that can lead to enhanced sensitivity to information that would otherwise have been determined to be redundant and irrelevant. This effect may extend beyond brief time periods to cause decreased long-term adaptation to stimuli as well, with the potential to create layers of sensory inputs that cannot be filtered out. Consequent overwhelming of sensory integration systems carries significant implications for understanding atypical processing and perception in autism. By better understanding these neurophysiologic mechanisms that may contribute to sensory overload or have downstream effects on social-emotional regulation, the field can move towards the development of effective therapeutics and behavioral strategies to improve autistic quality of life.

## Limitations

While this study included adults on the autism spectrum who had legal guardians and required a substantial amount of support, these participants were in the minority and the overall sample largely represents independent, verbally-fluent autistic adults. Given the simple, passive nature of this task, future work should aim to expand this research to individuals with autism who have higher support needs. Additionally, the consistency of timing within and between vibrotactile stimuli may have allowed for participants to anticipate the stimulus onset. While this approach allowed us to more precisely target prediction as a contributor to the pattern of responses observed, the extended ISI between stimulus sequences makes predicting the exact onset unlikely, and follow-up with randomized ISIs would allow for dissociation of prediction and re-sensitization of the first vibration in the sequence.

## DECLARATIONS

### List of Abbreviations

(ANOVA): Analysis of Variance
(ADHD): Attention-deficit/hyperactivity disorder
(ASD): Autism Spectrum Disorder
(EEG): Electroencephalography
(NT): Neurotypical

## Ethics approval and consent to participate

All aspects of the research conformed to the tenets outlined in the Declaration of Helsinki, with the exception that this study was not preregistered. The institutional review board of the University of Rochester, where the data collection took place, approved this study (STUDY00002036). All participants provided written informed consent.

## Consent for publication

Not applicable

## Author contributions

ELI and JJF conceived the study. ELI, JJF, and EGF designed the original experiment. ELI recruited and phenotyped the participants. ELI and ERL collected the data. ELI and ERL analyzed the data and created the illustrations and wrote the first draft of the paper, in close collaboration with JJF and EGF. JJF and EGF provided substantial editorial input and writing on subsequent drafts. All authors read the final draft and provided critical input.

## Acknowledgments

We thank Emily Knight, Erin Bojanek, Paige Nicklas, and Kathryn Toffolo for their support in participant recruitment and experimental design, and the entire team at the Frederick J. and Marion A. Schindler Cognitve Neurophysiology lab for their ongoing support of the work,

## Availability of data and material

The datasets used and/or analyzed during the current study are available upon request to the corresponding author.

## Clinical Trial

Not applicable.

## Literature cited

American Psychiatric Association. (2013). Diagnostic and statistical manual of mental disorders (5th ed.). American Psychiatric Publishing.

Andrade, G. N., Butler, J. S., Mercier, M. R., Molholm, S., & Foxe, J. J. (2015). Spatio-temporal dynamics of adaptation in the human visual system: a high-density electrical mapping study. Eur J Neurosci, 41(7), 925–939. 10.1111/ejn.12849

Andrade, G. N., Butler, J. S., Peters, G. A., Molholm, S., & Foxe, J. J. (2016). Atypical visual and somatosensory adaptation in schizophrenia-spectrum disorders. Translational psychiatry, 6(5), e804. 10.1038/tp.2016.63

Angel, R. W., Quick, W. M., Boylls, C. C., Weinrich, M., & Rodnitzky, R. L. (1985). Decrement of somatosensory evoked potentials during repetitive stimulation. Electroencephalogr Clin Neurophysiol, 60(4), 335–342. 10.1016/0013-4694(85)90007-0

Baranek, G. T., Foster, L. G., & Berkson, G. (1997). Tactile defensiveness and stereotyped behaviors. Am J Occup Ther, 51(2), 91–95. 10.5014/ajot.51.2.91

Barry, R. J., & James, A. L. (1988). Coding of stimulus parameters in autistic, retarded, and normal children: evidence for a two-factor theory of autism. Int J Psychophysiol, 6(2), 139–149. 10.1016/0167-8760(88)90045-1

Benjamini, Y., & Hochberg, Y. (1995). Controlling the False Discovery Rate: A Practical and Powerful Approach to Multiple Testing. Journal of the Royal Statistical Society. Series B (Methodological), 57(1), 289–300. http://www.jstor.org/stable/2346101

Blakemore, S. J., Tavassoli, T., Calò, S., Thomas, R. M., Catmur, C., Frith, U., & Haggard, P. (2006). Tactile sensitivity in Asperger syndrome. Brain Cogn, 61(1), 5–13. 10.1016/j.bandc.2005.12.013

Bruin, K. J., Kenemans, J. L., Verbaten, M. N., & Van der Heijden, A. H. (2000). Habituation: an event-related potential and dipole source analysis study. Int J Psychophysiol, 36(3), 199–209. 10.1016/s0167-8760(99)00114-2

Budd, T. W., Barry, R. J., Gordon, E., Rennie, C., & Michie, P. T. (1998). Decrement of the N1 auditory event-related potential with stimulus repetition: habituation vs. refractoriness. International Journal of Psychophysiology, 31(1), 51–68. 10.1016/S0167-8760(98)00040-3

Butler, J. S., Foxe, J. J., Fiebelkorn, I. C., Mercier, M. R., & Molholm, S. (2012). Multisensory representation of frequency across audition and touch: high density electrical mapping reveals early sensory-perceptual coupling. J Neurosci, 32(44), 15338–15344. 10.1523/jneurosci.1796-12.2012

Cascio, C., McGlone, F., Folger, S., Tannan, V., Baranek, G., Pelphrey, K. A., & Essick, G. (2008). Tactile perception in adults with autism: a multidimensional psychophysical study. J Autism Dev Disord, 38(1), 127–137. 10.1007/s10803-007-0370-8

Cauller, L. (1995). Layer I of primary sensory neocortex: where top-down converges upon bottom-up. Behav Brain Res, 71(1-2), 163–170. 10.1016/0166-4328(95)00032-1

Delorme, A., & Makeig, S. (2004). EEGLAB: an open source toolbox for analysis of single-trial EEG dynamics including independent component analysis. J Neurosci Methods, 134(1), 9–21. 10.1016/j.jneumeth.2003.10.009

Desmedt, J. E., Brunko, E., & Debecker, J. (1976). Maturation of the somatosensory evoked potentials in normal infants and children, with special reference to the early N1 component. Electroencephalography and clinical neurophysiology, 40(1), 43–58. 10.1016/0013-4694(76)90178-4

Dwyer, P., Williams, Z. J., Vukusic, S., Saron, C. D., & Rivera, S. M. (2023). Habituation of auditory responses in young autistic and neurotypical children. Autism Res, 16(10), 1903–1923. 10.1002/aur.3022

Eisenstein, E. M., Eisenstein, D., & Smith, J. C. (2001). The evolutionary significance of habituation and sensitization across phylogeny: A behavioral homeostasis model. Integrative Physiological & Behavioral Science, 36(4), 251–265. 10.1007/BF02688794

Espenhahn, S., Godfrey, K. J., Kaur, S., Ross, M., Nath, N., Dmitrieva, O., McMorris, C., Cortese, F., Wright, C., Murias, K., Dewey, D., Protzner, A. B., McCrimmon, A., Bray, S., & Harris, A. D. (2021). Tactile cortical responses and association with tactile reactivity in young children on the autism spectrum. Mol Autism, 12(1), 26. 10.1186/s13229-021-00435-9

Ferrer Knight, A., & Birtles, D. (2025). ‘I feel trapped in my safe clothes’: The impact of tactile hyper-sensitivity on autistic adults. Autism, 29(11), 2727–2740. 10.1177/13623613251366882

Foss-Feig, J. H., Heacock, J. L., & Cascio, C. J. (2012). TACTILE RESPONSIVENESS PATTERNS AND THEIR ASSOCIATION WITH CORE FEATURES IN AUTISM SPECTRUM DISORDERS. Res Autism Spectr Disord, 6(1), 337–344. 10.1016/j.rasd.2011.06.007

Francisco, A. A., Foxe, J. J., Horsthuis, D. J., & Molholm, S. (2022). Early visual processing and adaptation as markers of disease, not vulnerability: EEG evidence from 22q11.2 deletion syndrome, a population at high risk for schizophrenia. Schizophrenia, 8(1), 28. 10.1038/s41537-022-00240-0

Freedman, R., Adler, L. E., & Waldo, M. (1987). Gating of the Auditory Evoked Potential in Children and Adults. Psychophysiology, 24(2), 223–227. 10.1111/j.1469-8986.1987.tb00282.x

Gandhi, T. K., Tsourides, K., Singhal, N., Cardinaux, A., Jamal, W., Pantazis, D., Kjelgaard, M., & Sinha, P. (2021). Autonomic and Electrophysiological Evidence for Reduced Auditory Habituation in Autism. J Autism Dev Disord, 51(7), 2218–2228. 10.1007/s10803-020-04636-8

Green, S. A., Hernandez, L., Tottenham, N., Krasileva, K., Bookheimer, S. Y., & Dapretto, M. (2015). Neurobiology of Sensory Overresponsivity in Youth With Autism Spectrum Disorders. JAMA Psychiatry, 72(8), 778–786. 10.1001/jamapsychiatry.2015.0737

Groves, P. M., & Thompson, R. F. (1970). Habituation: A dual-process theory. Psychological Review, 77(5), 419–450. 10.1037/h0029810

Güçlü, B., Tanidir, C., Mukaddes, N. M., & Unal, F. (2007). Tactile sensitivity of normal and autistic children. Somatosens Mot Res, 24(1-2), 21–33. 10.1080/08990220601179418

Guiraud, J. A., Kushnerenko, E., Tomalski, P., Davies, K., Ribeiro, H., & Johnson, M. H. (2011). Differential habituation to repeated sounds in infants at high risk for autism. Neuroreport, 22(16), 845–849. 10.1097/WNR.0b013e32834c0bec

Harris, J. D. (1943). Habituatory response decrement in the intact organism. Psychological Bulletin, 40(6), 385–422. 10.1037/h0053918

Hashimoto, T., Tayama, M., & Miyao, M. (1986). Short latency somatosensory evoked potentials in children with autism. Brain Dev, 8(4), 428–432. 10.1016/s0387-7604(86)80065-1

Isenstein, E. L., Freedman, E. G., Molholm, S., & Foxe, J. J. (2024). Somatosensory temporal sensitivity in adults on the autism spectrum: A high-density electrophysiological mapping study using the mismatch negativity (MMN) sensory memory paradigm. Autism Res. 10.1002/aur.3186

Isenstein, E. L., Freedman, E. G., Rico, G. A., Brown, Z., Tadin, D., & Foxe, J. J. (2025). Adults on the autism spectrum differ from neurotypical peers when self-generating but not passively-experiencing somatosensation: a high-density electrophysiological (EEG) mapping and virtual reality study. Neuroimage, 311, 121215. 10.1016/j.neuroimage.2025.121215

Isenstein, E. L., Grosman, H. E., Guillory, S. B., Zhang, Y., Barkley, S., McLaughlin, C. S., Levy, T., Halpern, D., Siper, P. M., Buxbaum, J. D., Kolevzon, A., & Foss-Feig, J. H. (2022). Neural Markers of Auditory Response and Habituation in Phelan-McDermid Syndrome [Original Research]. Frontiers in Neuroscience, 16. 10.3389/fnins.2022.815933

Jamal, W., Cardinaux, A., Haskins, A. J., Kjelgaard, M., & Sinha, P. (2021). Reduced Sensory Habituation in Autism and Its Correlation with Behavioral Measures. J Autism Dev Disord, 51(9), 3153–3164. 10.1007/s10803-020-04780-1

Jamioł-Milc, D., Bloch, M., Liput, M., Stachowska, L., & Skonieczna-Żydecka, K. (2021). Tactile Processing and Quality of Sleep in Autism Spectrum Disorders. Brain Sci, 11(3). 10.3390/brainsci11030362

Kurz, M. J., Wiesman, A. I., Coolidge, N. M., & Wilson, T. W. (2017). Children with Cerebral Palsy Hyper-Gate Somatosensory Stimulations of the Foot. Cerebral Cortex, 28(7), 2431–2438. 10.1093/cercor/bhx144

Lawson, R. P., Aylward, J., White, S., & Rees, G. (2015). A striking reduction of simple loudness adaptation in autism. Sci Rep, 5, 16157. 10.1038/srep16157

Liu, Y., Wang, W., Xu, W., Cheng, Q., & Ming, D. (2021). Quantifying the Generation Process of Multi-Level Tactile Sensations via ERP Component Investigation. International Journal of Neural Systems, 31(12), 2150049. 10.1142/s0129065721500490

Mammen, M. A., Moore, G. A., Scaramella, L. V., Reiss, D., Ganiban, J. M., Shaw, D. S., Leve, L. D., & Neiderhiser, J. M. (2015). INFANT AVOIDANCE DURING A TACTILE TASK PREDICTS AUTISM SPECTRUM BEHAVIORS IN TODDLERHOOD. Infant Ment Health J, 36(6), 575–587. 10.1002/imhj.21539

Mancini, F., Pepe, A., Bernacchia, A., Stefano, G. D., Mouraux, A., & Iannetti, G. D. (2018). Characterizing the Short-Term Habituation of Event-Related Evoked Potentials. eneuro, 5(5), ENEURO.0014-0018.2018. 10.1523/eneuro.0014-18.2018

McKernan, E. P., Wu, Y., & Russo, N. (2020). Sensory Overresponsivity as a Predictor of Amplitude Discrimination Performance in Youth with ASD. Journal of Autism and Developmental Disorders, 50(9), 3140–3148. 10.1007/s10803-019-04013-0

Megela, A. L., & Teyler, T. J. (1979). Habituation and the human evoked potential. Journal of Comparative and Physiological Psychology, 93(6), 1154–1170. 10.1037/h0077630

Mikkelsen, M., Wodka, E. L., Mostofsky, S. H., & Puts, N. A. J. (2018). Autism spectrum disorder in the scope of tactile processing. Dev Cogn Neurosci, 29, 140–150. 10.1016/j.dcn.2016.12.005

Montoya, P., Sitges, C., García-Herrera, M., Rodríguez-Cotes, A., Izquierdo, R., Truyols, M., & Collado, D. (2006). Reduced brain habituation to somatosensory stimulation in patients with fibromyalgia. Arthritis Rheum, 54(6), 1995–2003. 10.1002/art.21910

Muenssinger, J., Stingl, K., Matuz, T., Binder, G., Ehehalt, S., & Preissl, H. (2013). Auditory habituation to simple tones: reduced evidence for habituation in children compared to adults [Original Research]. Frontiers in Human Neuroscience, 7. 10.3389/fnhum.2013.00377

Müller, J. R., Metha, A. B., Krauskopf, J., & Lennie, P. (1999). Rapid adaptation in visual cortex to the structure of images. Science, 285(5432), 1405–1408. 10.1126/science.285.5432.1405

Nakajima, Y., & Imamura, N. (2000). Probability and Interstimulus Interval Effects on The N140 and the P300 Components of Somatosensory Erps. International Journal of Neuroscience, 104(1), 75–91. 10.3109/00207450009035010

O’Riordan, M., & Passetti, F. (2006). Discrimination in autism within different sensory modalities. J Autism Dev Disord, 36(5), 665–675. 10.1007/s10803-006-0106-1

Pellicano, E., Jeffery, L., Burr, D., & Rhodes, G. (2007). Abnormal Adaptive Face-Coding Mechanisms in Children with Autism Spectrum Disorder. Current Biology, 17(17), 1508–1512. 10.1016/j.cub.2007.07.065

Podoly, T. Y., & Ben-Sasson, A. (2020). Sensory Habituation as a Shared Mechanism for Sensory Over-Responsivity and Obsessive-Compulsive Symptoms. Front Integr Neurosci, 14, 17. 10.3389/fnint.2020.00017

Puts, N. A., Wodka, E. L., Tommerdahl, M., Mostofsky, S. H., & Edden, R. A. (2014). Impaired tactile processing in children with autism spectrum disorder. J Neurophysiol, 111(9), 1803–1811. 10.1152/jn.00890.2013

Puts, N. A. J., Harris, A. D., Crocetti, D., Nettles, C., Singer, H. S., Tommerdahl, M., Edden, R. A. E., & Mostofsky, S. H. (2015). Reduced GABAergic inhibition and abnormal sensory symptoms in children with Tourette syndrome. Journal of Neurophysiology, 114(2), 808–817. 10.1152/jn.00060.2015

Puts, N. A. J., Wodka, E. L., Harris, A. D., Crocetti, D., Tommerdahl, M., Mostofsky, S. H., & Edden, R. A. E. (2017). Reduced GABA and altered somatosensory function in children with autism spectrum disorder. Autism Res, 10(4), 608–619. 10.1002/aur.1691

Rankin, C. H., Abrams, T., Barry, R. J., Bhatnagar, S., Clayton, D. F., Colombo, J., Coppola, G., Geyer, M. A., Glanzman, D. L., Marsland, S., McSweeney, F. K., Wilson, D. A., Wu, C.-F., & Thompson, R. F. (2009). Habituation revisited: An updated and revised description of the behavioral characteristics of habituation. Neurobiology of Learning and Memory, 92(2), 135–138. 10.1016/j.nlm.2008.09.012

Ren, R., Yu, Y., Tang, X., Suzumura, S., Ejima, Y., Wu, J., & Yang, J. (2025). Electrophysiological evidence for the effect of tactile temporal prediction. Neuropsychologia, 210, 109095. 10.1016/j.neuropsychologia.2025.109095

Rogers, S. J., Hepburn, S., & Wehner, E. (2003). Parent reports of sensory symptoms in toddlers with autism and those with other developmental disorders. J Autism Dev Disord, 33(6), 631–642. 10.1023/b:jadd.0000006000.38991.a7

Rosburg, T., Zimmerer, K., & Huonker, R. (2010). Short-term habituation of auditory evoked potential and neuromagnetic field components in dependence of the interstimulus interval. Exp Brain Res, 205(4), 559–570. 10.1007/s00221-010-2391-3

Russo, N., Foxe, J. J., Brandwein, A. B., Altschuler, T., Gomes, H., & Molholm, S. (2010). Multisensory processing in children with autism: high-density electrical mapping of auditory-somatosensory integration. Autism Res, 3(5), 253–267. 10.1002/aur.152

Sapey-Triomphe, L. A., Sanchez, G., Hénaff, M. A., Sonié, S., Schmitz, C., & Mattout, J. (2023). Disentangling sensory precision and prior expectation of change in autism during tactile discrimination. NPJ Sci Learn, 8(1), 54. 10.1038/s41539-023-00207-5

Schröder, P., Nierhaus, T., & Blankenburg, F. (2021). Dissociating Perceptual Awareness and Postperceptual Processing: The P300 Is Not a Reliable Marker of Somatosensory Target Detection. J Neurosci, 41(21), 4686–4696. 10.1523/jneurosci.2950-20.2021

Schwartz, S., Vuilleumier, P., Hutton, C., Maravita, A., Dolan, R. J., & Driver, J. (2004). Attentional Load and Sensory Competition in Human Vision: Modulation of fMRI Responses by Load at Fixation during Task-irrelevant Stimulation in the Peripheral Visual Field. Cerebral Cortex, 15(6), 770–786. 10.1093/cercor/bhh178

Sinha, P., Kjelgaard, M. M., Gandhi, T. K., Tsourides, K., Cardinaux, A. L., Pantazis, D., Diamond, S. P., & Held, R. M. (2014). Autism as a disorder of prediction. Proc Natl Acad Sci U S A, 111(42), 15220–15225. 10.1073/pnas.1416797111

Tarkka, I. M., Treede, R. D., & Bromm, B. (1992). Sensory and movement-related cortical potentials in nociceptive and auditory reaction time tasks. Acta Neurol Scand, 86(4), 359–364. 10.1111/j.1600-0404.1992.tb05101.x

Tavassoli, T., Bellesheim, K., Tommerdahl, M., Holden, J. M., Kolevzon, A., & Buxbaum, J. D. (2016). Altered tactile processing in children with autism spectrum disorder. Autism Res, 9(6), 616–620. 10.1002/aur.1563

Thompson, R. F., & Spencer, W. A. (1966). Habituation: A model phenomenon for the study of neuronal substrates of behavior. Psychological Review, 73(1), 16–43. 10.1037/h0022681

Uppal, N., Foxe, J. J., Butler, J. S., Acluche, F., & Molholm, S. (2016). The neural dynamics of somatosensory processing and adaptation across childhood: a high-density electrical mapping study. J Neurophysiol, 115(3), 1605–1619. 10.1152/jn.01059.2015

van der Miesen, M. M., Joosten, E. A., Kaas, A. L., Linden, D. E. J., Peters, J. C., & Vossen, C. J. (2024). Habituation to pain: self-report, electroencephalography, and functional magnetic resonance imaging in healthy individuals. A scoping review and future recommendations. PAIN, 165(3), 500–522. 10.1097/j.pain.0000000000003052

Villalonga, M. B., Sussman, R. F., & Sekuler, R. (2021). Perceptual timing precision with vibrotactile, auditory, and multisensory stimuli. Atten Percept Psychophys, 83(5), 2267–2280. 10.3758/s13414-021-02254-9

Wang, A. L., Mouraux, A., Liang, M., & Iannetti, G. D. (2008). The enhancement of the N1 wave elicited by sensory stimuli presented at very short inter-stimulus intervals is a general feature across sensory systems. PLoS One, 3(12), e3929. 10.1371/journal.pone.0003929

Webb, S. J., Jones, E. J., Merkle, K., Namkung, J., Toth, K., Greenson, J., Murias, M., & Dawson, G. (2010). Toddlers with elevated autism symptoms show slowed habituation to faces. Child Neuropsychol, 16(3), 255–278. 10.1080/09297041003601454

World Medical Association Declaration of Helsinki: ethical principles for medical research involving human subjects. (2013). Jama, 310(20), 2191–2194. 10.1001/jama.2013.281053

Yamaguchi, S., & Knight, R. T. (1991). P300 generation by novel somatosensory stimuli. Electroencephalogr Clin Neurophysiol, 78(1), 50–55. 10.1016/0013-4694(91)90018-y

Zanini, S., Martucci, L., Del Piero, I., & Restuccia, D. (2016). Cortical hyper-excitability in healthy children: evidence from habituation and recovery cycle phenomena of somatosensory evoked potentials. Developmental Medicine & Child Neurology, 58(8), 855–860. 10.1111/dmcn.13072

